# Beta diversity patterns derived from island biogeography theory

**DOI:** 10.1101/292490

**Authors:** Muyang Lu, David Vasseur, Walter Jetz

## Abstract

The Theory of Island Biogeography (TIB) has been successful in predicting alpha diversity patterns such as species-area relationships and species-abundance distributions. Although beta diversity (i.e. the dissimilarity of community composition) has long been recognized as an important element of the TIB and is crucial for understanding community assembly processes, it has never been formally incorporated into the theory. Here we derive theoretical predictions for the expected pairwise beta diversity values under a species-level neutral scenario where all species have equal colonization and extinction rates. We test these predictions for the avian community composition of 42 islands (and 93 species) in the Thousand Island Lake, China. We find that alpha diversity patterns alone do not distinguish a species-level neutral model from a non-neutral model. In contrast, beta diversity patterns clearly reject a species-level neutral model. We suggest that the presented theoretical integration beta diversity offers a powerful path for testing the presence of neutral processes in ecology and biogeography.

## INTRODUCTION

Island biogeography is undergoing a renaissance (Warren et al. 2015; Santos et al. 2016; Patiño et al. 2017), at the heart of which lies the equilibrium theory of island biogeography (TIB) formulated by MacArthur and Wilson 50 years ago (1967). The TIB is most celebrated for elegantly linking species richness with colonization and extinction rates, which in turn are influenced by island area and distance to the mainland. Equilibrium TIB makes two key predictions: 1) equilibrium species richness depends on colonization and extinction rates; 2) community composition of an island is undergoing constant turnover when at equilibrium. While early empirical studies have focused on testing the equilibrium and temporal turnover predictions (Diamond 1969; Simberloff 1969; Simberloff and Wilson 1969, 1970; Diamond and May 1977), recent interest in TIB is stimulated by the success of Hubbell’s unified neutral theory (a descendant of the TIB; Hubbell 2001), the advance of food web theories (Holt 2009), the developments of phylogenetic analyses (Valente et al. 2015; Lim and Marshall 2017), and the accumulation of island datasets (Weigelt et al. 2013, 2016). Motivated by the desire for a general island biogeography theory, more and more complexities have been added to the original framework, including allometric scaling (Jacquet et al. 2017), trophic interactions (Gravel et al. 2011; Cazelles et al. 2016), speciation (Chen and He 2009; Rosindell and Phillimore 2011; Rosindell and Harmon 2013), habitat heterogeneity (Kadmon and Allouche 2007), and island ontogenies (Whittaker et al. 2008; Borregaard et al. 2016). While the complexity of the theory has increased, examinations to date have focused on alpha diversity patterns such as species richness (MacArthur and Wilson 1967), species-abundance relationships (Rosindell and Phillimore 2011; Rosindell and Harmon 2013; Kessler and Shnerb 2015), functional diversity (Jacquet et al. 2017), and endemism (Chen and He 2009).

One particular omission in the theory and tests of the TIB to date is beta diversity, i.e. the compositional dissimilarities among communities, a widely used metric in biodiversity studies (Leprieur et al. 2011; Stegen et al. 2013; Segre et al. 2014; Si et al. 2016). The integration of beta diversity patterns into the TIB has the potential to unlock a range of uses in community assembly studies and the development of a unified meta-community theory (Leibold et al. 2004).

The concept of beta diversity was first introduced by Whittaker (1960, 1965) and was quickly linked to an array of important mechanisms in community assembly: biotic interaction (Graham and Fine 2008), environment filtering (Veech and Crist 2007; Buckley and Jetz 2008), dispersal limitation (Ojima and Jiang 2017; Wu et al. 2017), and historical contingency (Fukami and Nakajima 2011). However, despite the widespread recognition of the beta diversity concept, there is little consensus about the best way to measure it (Tuomisto 2010a, 2010b; Anderson et al. 2011).

Depending on the purpose of a study, beta diversity could be measured as turnover (directional) or variation (non-directional) of compositional similarity (Nekola and White 1999; Legendre and De Cáceres 2013); based on pairwise or multiple-sites comparisons (Hui and McGeoch 2014; Arita 2017; Latombe et al. 2017; Marion et al. 2017); decomposed in additive or multiplicative framework (with respect to alpha and gamma diversity; Jost 2007; Cabral *et al.* 2014); calculated by incidence (presence-absence) or abundance data (Chao et al. 2016). Even for the commonly used pairwise beta diversity, there is a heated debate about what is the best way to partition it into richness difference and replacement (or nestedness and turnover) components (Baselga 2010; Podani and Schmera 2011; Legendre 2014; Baselga and Leprieur 2015). It is generally agreed that different measures quantify different aspects of communities and that their use should be guided by the question asked (Anderson et al. 2011; Legendre 2014). Although beta-diversity measures have been increasingly synthesized under more general frameworks such as Hill’s number and variance of community matrix (Chao et al. 2016), only a few of them have been examined analytically from process-based theories (Chave and Leigh 2002; Condit et al. 2002; Zillio et al. 2005; Connolly et al. 2017). Moreover, theoretical considerations of beta-diversity patterns to date have been abundance-based and hence limited to more restricted datasets, such as vegetation plots (Chave and Leigh 2002; Condit et al. 2002; Zillio et al. 2005). In summary, the link between beta diversity patterns and fundamental processes such as colonization and extinction in the TIB remain ill developed.

In this study, we set out to address this shortcoming and provide a theoretical and example empirical integration of beta diversity into TIB. Specifically, we present the first theoretical predictions for the expected pairwise beta diversity patterns for both Jaccard and Sørensen indices in the classic MacArthur and Wilson framework. This framework constitutes in essence a species-level neutral model (as opposed to the individual-level neutral theory; Hubbell 2009; Rosindell and Phillimore 2011). The species-level neutral model assumes that all species have the same colonization and extinction rates, while the individual-level neutral model assumes that all individuals have the same colonization and extinction rates, which allows species-level asymmetry to arise because of different abundance (Rosindell and Harmon 2013). We first derived the probability mass function and expected value of pairwise beta diversity in a general case where the equilibrium assumption is not required. We then examined our results under the classic MacArthur-Wilson equilibrium framework. Finally, we test the derived predictions for a dataset of 93 avian species occurring across 42 islands of the Thousand Island Lake Region in China (Appendix, Wang *et al.* 2010).

## MATERIAL AND METHODS

To derive expected beta diversity, we need to consider the joint statistical distribution of multiple islands. We here focus on pairwise community comparisons, because average pairwise dissimilarity is shown to be the only unbiased and consistent estimator when applied to empirical data (Marion et al. 2017). We also calculated the expected value of the partitioned components of pairwise beta dissimilarity for both Baselga and Podani families (Baselga 2010; Podani and Schmera 2011). To test our predictions, we applied an “Incidence Function” approach pioneered by Diamond (1975) and used by Connor and Simberloff (1978, 1979) during early debates of null models.

### Probability mass function of pairwise beta diversity

All pairwise beta diversity indices share the same probability mass function to describe island community composition because they are calculated from four quantities: the total number of species of two islands *N* (regional species richness), the number of species shared by both islands *k*, the number of species unique to the first island *i*, and the number of species unique to the second island *j* (thus *N* = *k* + *i + j*). Then the number of species present on the first island is *i*+*k*, and the number of species present on the second island is *j*+*k.*

Denoting the size of the mainland species pool as *M*, the probability mass function of pairwise beta diversity, conditioning on *N*, *k* and *i,* can be derived as follows: let the occurrence probability of the *i*^th^ species on the first island be *p*, and the *j*^th^ species on the second island be *q*. Furthermore, assuming that the occurrence probabilities are properties of the island rather than of species, the probability of obtaining exactly *i* unique species and *k* shared species from a regional pool of size *M* is given by *p*^*k+i*^(1-*p*)^*M*-(*k*+*i*)^. Similarly, the probability of obtaining *j* unique and *k* shared species on the second island is *q*^*k+j*^(1-*q*)^*M*-(*k*+*j*)^, which yields the probability of obtaining the regional community specified by *i, j, k* and *M*, as *p*^*k*+*i*^(1-*p*)^*M*-(*k*+*i*)^*q^k^*^+*j*^(1-*q*)^*M*-(*k*+*j*)^.

Assuming that species are neutral with respect to their occurrence probabilities within an island, we then count how many different combinations of island communities could be obtained with regional richness *N*, *k* shared species, and *i* unique species to the first island (the number of unique species to the second island is then fixed as *k-i*). There are exactly 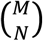 ways of choosing *N* out of *M* species to be present in either or both islands from the mainland species pool. Similarly, there are 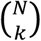 ways to choose *k* out of *N* species to be shared by both islands and for the unique *N-k* species, there are 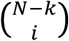 ways of assigning *i* species to the first island. After simplification, the probability of pairwise beta diversity conditioning on the total number of species of both islands *N*, the number of species shared by both islands *k*, and the number of species unique to the first island *i* is:

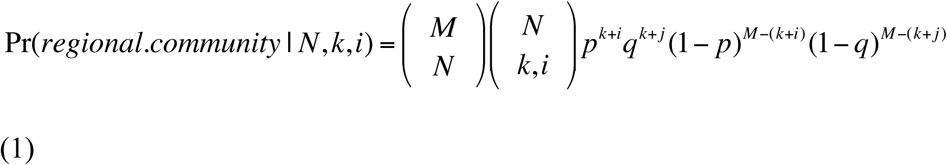

Because beta diversity measures (e.g. Jaccard and Sørensen indices) are usually undefined when *N* = 0, the expected pairwise beta diversity should be normalized by the term 1-(1-*p*)^*M*^(1-*q*)^*M*^ to exclude “double-absence” scenarios (Anderson et al. 2011). Only the expected Jaccard dissimilarity (Jaccard dissimilarity is defined as 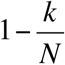) has a simple analytical form that is independent of the size of mainland species pool *M* (see Appendix S1 in Supporting Information for derivation; we do not show the analytical form of the expected Sørensen dissimilarity because the specific form depends on the size of mainland species pool):

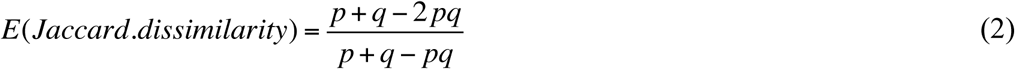

It has a form that is similar to the expected pairwise Jaccard derived by Chase *et al.* (2011) but allows two islands to have different occurrence probabilities. *p* and *q* could also be interpreted as probabilities of the same island at different times, and in that case (2) becomes the expected temporal Jaccard dissimilarity. This quantity does not require any equilibrium assumptions of alpha diversity and hence could be seen as a generalized version of the apparent turnover derived by Diamond and May (1977) which is essentially a pairwise Sørensen dissimilarity index in equilibrium (Morrison 2017).

We also calculated the expected pairwise beta diversity for Jaccard and Sørensen families as well as their partitioned components (for detailed formulas of the Baselga and Podani families see Baselga & Leprieur 2015) conditioning on both islands having species (*i+k* >0 and *j+k* > 0) because empty islands are sometimes excluded from analysis (Wang et al. 2016). In Baselga’s family, pairwise dissimilarity is partitioned into the nestedness component and the turnover component. The nestedness component measures the extent to which species in a smaller community are a subset of species in a larger community, while the turnover component measures how much of the dissimilarity is caused by species replacement (Baselga 2010). The turnover component and the nestedness component in Baselga’s family, respectively, correspond to the replacement component and the richness difference component in Podani’s family (Legendre 2014).

### Island biogeography theory

Following the stochastic version of the TIB (MacArthur and Wilson 1967), the occurrence probability of a species on an island is a function of colonization rate *c* and extinction rate *e*:

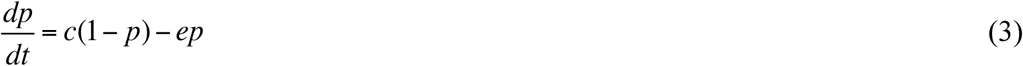

When occurrence probability of a species is at stochastic equilibrium (stationary distribution), the occurrence probability is:

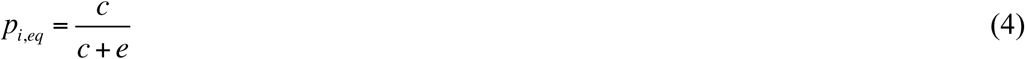

Let 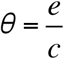, which is the relative extinction rates, equation (4) becomes:

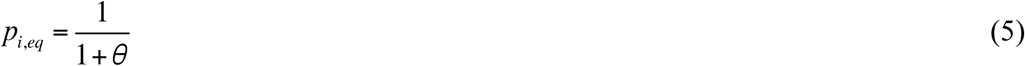

Substitute 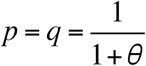 into equation (2), the expected Jaccard dissimilarity when two islands have equal colonization and extinction probabilities is:

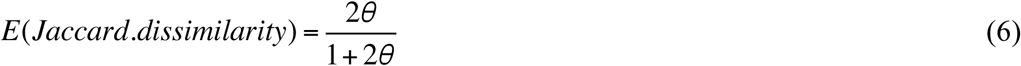

To take into account asymmetry between islands (e.g. the effect of area, isolation or habitat types), let 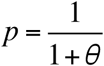 and 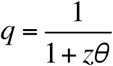, so that the relative extinction rate on the second island is *z* times the relative extinction rate on the first island. Equation (2) becomes:

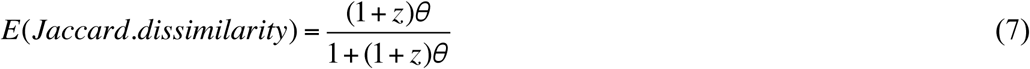

### Empirical tests Data

We use a published dataset of avian community composition for 93 birds and 42 islands in The Thousand Island Lake, China to test our predictions (see Appendix in Wang *et al.* 2010). The region (29°22˝ – 29°50˝ N, 118°34˝– 119°15˝ E) consists of an inundated lake with more than 1000 islands created by dam constructions in 1959. Because the islands were formed recently, there was no in situ speciation in this region. The relative small area of the region (573 km^2^) ensures that the islands are accessible to most of the bird species (Si et al. 2016). Bird occupancies from 2006 to 2009 on 42 islands were documented using line-transect method. Island variables measured in the dataset include area, distance to mainland, habitat diversity and plant species richness.

### Incidence function and statistical analysis

Because extinction rates and colonization rates are difficult to measure directly, an alternative way is to use an “Incidence function” approach to estimate parameters from a snapshot of occupancy patterns (Diamond 1975; Hanski 2009). To test the predictions of island biogeography theory, equation (4) is modeled as a function of isolation and area. Parameters are fitted by maximum likelihood with binomial distribution. Three neutral models and one non-neutral model are examined in this study:

#### Neutral model 1: Inverse ratio

Colonization rates are modeled as an inverse ratio function of isolation: 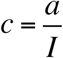, and extinction rates are modeled as an inverse ratio of area: 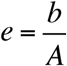. *a* and *b* are fitted parameters.

#### Neutral model 2: Exponential

Colonization rates are modeled as an exponential function of isolation:

*c* = exp(—*aI*) and extinction rates are modeled as an exponential function of area:

*e* = exp(—*bA*)*a* and *b* are fitted parameters. Substitute *c* = exp(—*aI*) and

*e* = exp(—*bA*) into equation (4), the occurrence probability becomes 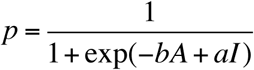

Thus the exponential neutral model is equivalent to a logistic regression without intercepts, which could be directly fitted by ‘glm’ in R with binomial distribution and logit link.

#### Neutral model 3: GLM

This model adds habitat diversity and plant richness to the predictors in addition to area and isolation in Neutral Model 2. AIC is used to select the best model. Area and isolation are log-transformed to ensure better linearity.

Non-neutral model: Aggregate species-level GLM

To take into account species-level non-neutrality, species identity is included as a fixed effect categorical variable in the GLM, which allows each species to have a different intercept (baseline occurrence probability) but share the same response to island area, isolation, plant richness and habitat types. This is essentially a stacked species distribution modeling approach (Calabrese et al. 2014; Ko et al. 2016), which increases the number of parameters by the number of species minus one (92 parameters in this case). This procedure aims to capture the observed presence-absence variation among species but does not pin down the causes of species-level non-neutrality such as traits and abundance differences. More realistic assumptions of non-neutrality such as different responses to area and isolation could be made but are not the main focus of this paper. AUC and AIC values are calculated to compare the overall performance of the models. AUC is calculated with a Mann-Whitney U statistic using R package ‘PresenceAbsence’. AIC is calculated from the best-fitted likelihood function. ΔAIC are calculated by subtracting the AIC of the null model (GLM with only one intercept) from the AIC of the fitted model.

### Predictions of alpha diversity and beta diversity

The predicted species richness of each island is given by the summed fitted occurrence probabilities of all species as used in conventional stacked species distribution modeling (Calabrese et al. 2014; Ko et al. 2016). While predicted pairwise Jaccard dissimilarity can be calculated analytically from equation (2), the partitioned nestedness and turnover components can only be estimated by simulations (we only present the results of Baselga’s family because its partitioned components are independent with each other when gamma diversity is fixed; Baselga & Leprieur 2015). We therefore estimate predicted pairwise beta diversity as the mean of 1000 community matrices simulated from the fitted occurrence probabilities. Observed values are regressed against predicted values using ordinary least square. *R*^2^ is used as a measure of predictive power for alpha and beta diversity patterns. If the model predicts the observed patterns well, the fitted regression line should be close to the 1:1 ratio line when observed values are plotted against predicted values. All statistical analyses are performed in R 3.3.2.

## RESULTS

### Symmetric islands (same colonization and extinction rates for both islands)

The expected Jaccard dissimilarity conditional on both islands having non-zero species richness increases monotonically with relative extinction rate *θ* and the size (number of species) of the mainland species pool *M*. As *M* increases, the expected Jaccard dissimilarity converges to 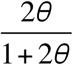 (Fig. 1a). When decomposed into turnover and nestedness (or replacement and richness difference) components, contrasting patterns are observed: while the turnover component and the replacement component increase monotonically with *θ* and *M* (Fig. 1b, e), the nestedness component and richness difference component are both unimodal functions of *θ* with maximum values less than 0.3 (Fig. 1c, f). The maximum nestedness decreases as *M* increases (Fig. 1c), while the maximum richness difference changes little with the increase of *M* (Fig. 1f). The relative importance of turnover increases monotonically with *θ* and *M* (Fig. 1d). In a special case of 2 mainland species, the ratio of expected turnover and expected nestedness equals *θ* (Fig. 1d). The Sørensen family indices have similar quantitative behaviors as the Jaccard family indices (see Fig. S1 in Appendix).

**Figure 1.**
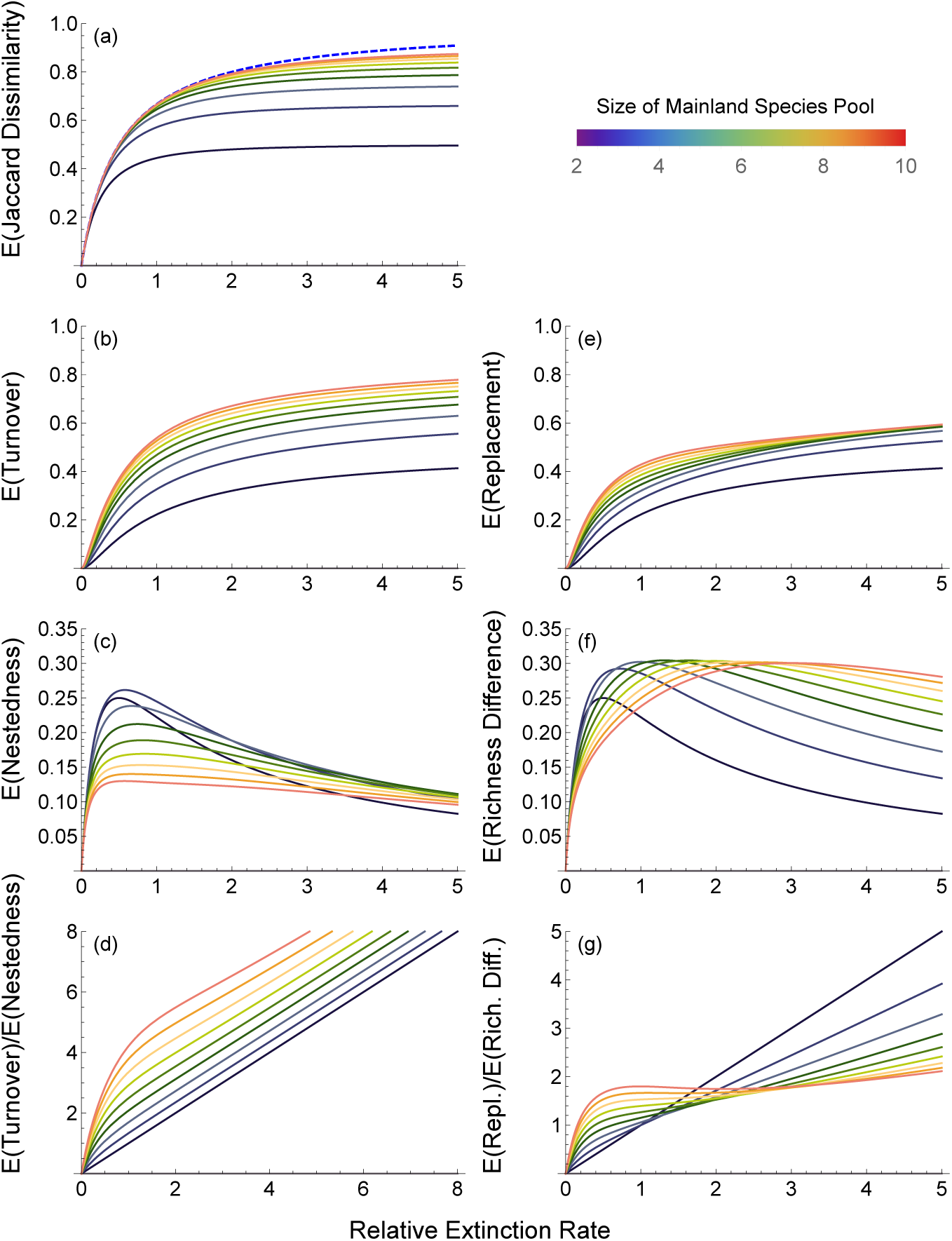
Expected pairwise beta diversity of two identical islands (same colonization and extinction rates), conditioning on both islands having species (*i* > 0, *j* > 0). Results are derived from the joint presence-absence distribution of two islands and shown for different mainland pool sizes. Blue dashed line in panel (a) is the analytical solution 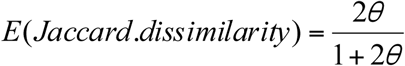 for the case of at least one island having species (*N >* 0). Relative extinction rate is the ratio of extinction rate and colonization rate.

### Asymmetric islands (different colonization and extinction rates for two islands)

The expected Jaccard dissimilarity conditional on both islands having non-zero species richness converges to 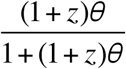 as *M* increses. But the deviation from 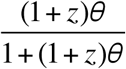 also increses with island asymmetry *z* (Fig. 2a, e, i). The turnover component increases monotonically with *θ* and *M*, and decreases with *z* (Fig. 2b, f, j). The nestedness component decreases with *M* when *z* is small (Fig. 2c), but this relationship with *M* is reversed when *z* gets larger (Fig. 2g, k). The ratio of expected turnover and expected nestedness also increases with *M* when *z* is small (Fig. 2d), and decreases with *M* when *z* is large (Fig. 2h, l).

**Figure 2.**
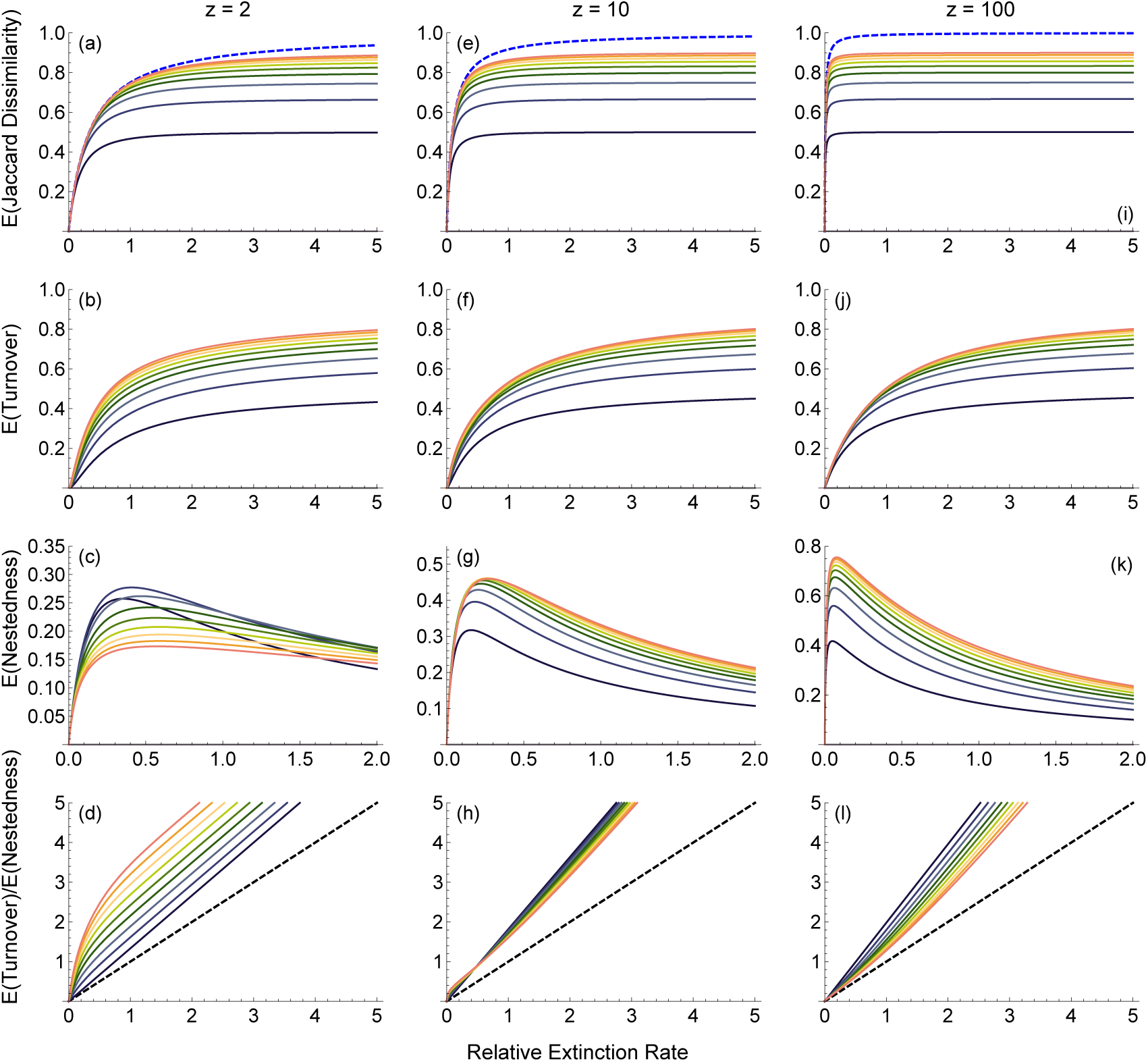
Expected pairwise Jaccard beta diversities of two islands differing in colonization and/or extinction rates, conditioning on both islands having species (*i* > 0, *j* > 0). The blue dashed line in panel (a), (e) and (i) are the analytical solutions of 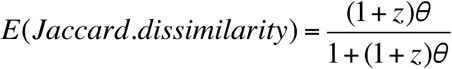, conditional on at least one island having nonzero species. Expectations are provided for different levels of z, i.e. the number of times relative extinction rate on the second island exceeds that on the first. The black dashed lines in panel (d), (h) and (l) represent a 1:1 ratio. Different lines in a graph represent different mainland pool sizes (for color legend see Fig. 1).

### Empirical test

We tested the above predictions for an inland lake island system using 93 bird species and 42 islands. We used an incidence function approach to fit the observed occupancy patterns and calculated predicted alpha and beta diversity patterns from the fitted models. The non-neutral GLM has the lowest AIC and highest AUC in all models (Table 1). Among three neutral models, the neutral GLM model has the lowest AIC and the highest AUC. All models are better at predicting alpha diversity patterns than at predicting beta diversity patterns (Fig. 3). The neutral exponential model and GLM systematically underestimate the nestedness component, but overestimate the turnover component (Fig. 3f, j). In contrast, the neutral inverse ratio model systematically underestimates all observed pairwise beta diversity at the lower range of the predictions, yet it overestimates pairwise beta diversity at the higher range of the predictions (Fig. 3a-d). Both the neutral GLM and the non-neutral GLM successfully predict the observed alpha diversity pattern (Fig. 3i, m), but only the non-neutral GLM successfully predicts the observed beta diversity patterns (Fig. 3n-p).

**Figure 3.**
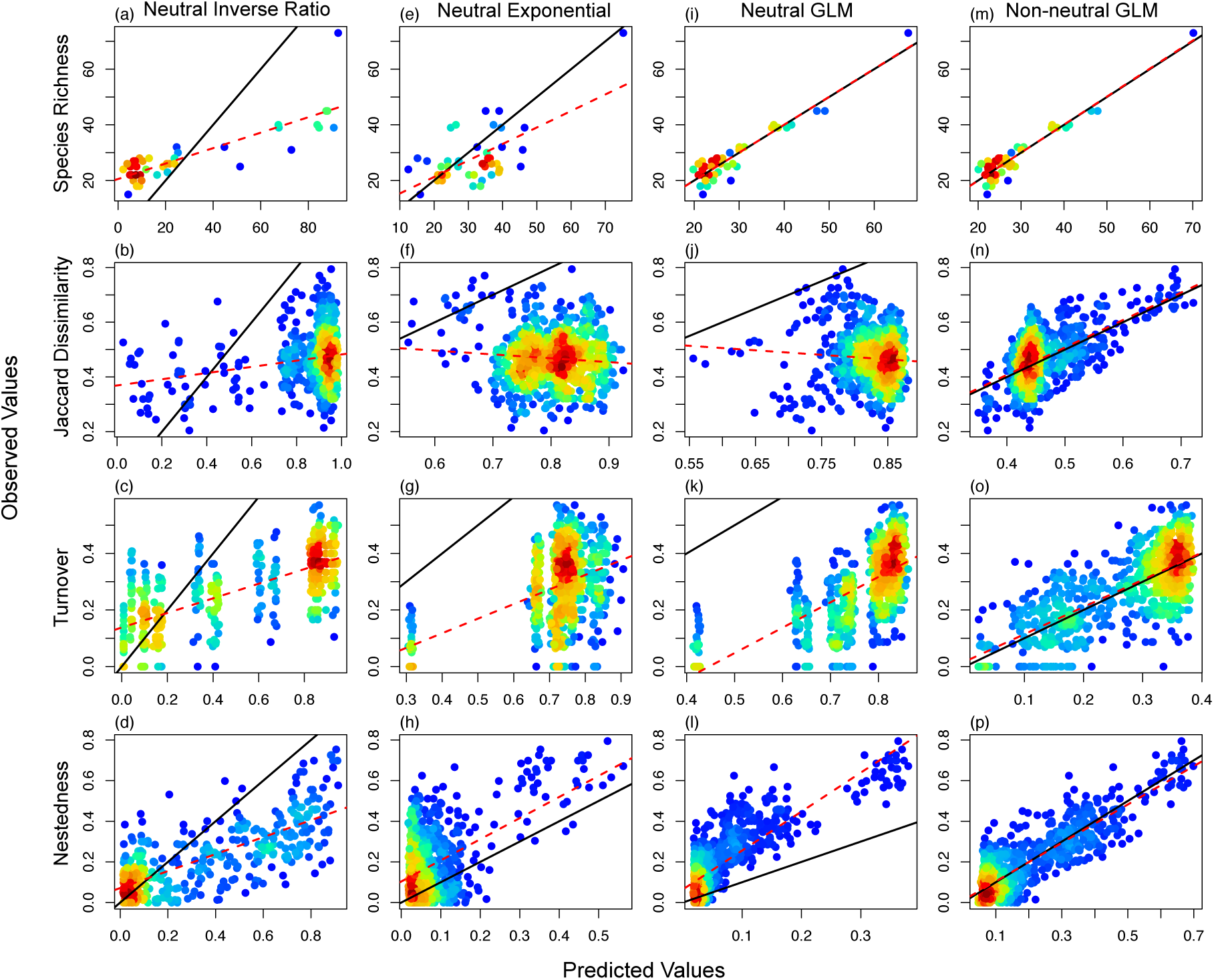
Empirical evaluation of the theoretically fitted alpha and beta diversity patterns for the Thousand Islands Lake bird dataset. (a) - (l) are predictions of three neutral models, (m) - (p) are predictions of a non-neutral model. Solid black lines represent a 1:1 relationship and dashed red lines are fits of ordinary linear regressions, Point densities from high to low are color-coded from red to blue.

**Table 1.** Models fitted to the presence-absence patterns of 93 bird species in 42 islands in the Thousand Island Lake, China.

| Model | $\Delta AIC$ | AUC | Number of<br>parameters | $R^2$ | | |
| --- | --- | --- | --- | --- | --- | --- |
|  |  |  |  | Richness | Dissimilarity | Turno |
| non-neutral.GLM | -2627 | 0.95 | 97 | 0.91 | 0.37 | 0.53 |
| neutral.GLM | -172 | 0.61 | 5 | 0.90 | 0.01 | 0.45 |
| neutral.exponential | -8 | 0.57 | 2 | 0.41 | 0.01 | 0.17 |
| neutral.inverse ratio | 1587 | 0.60 | 1 | 0.69 | 0.04 | 0.46 |
| Total observation = 3906 |  |  |  |  |  |  |

Note: For the neutral exponential model: $c = \exp(-aI)$ and $e = \exp(-bA)$ . For the neutral inverse ratio model: $c = \frac{a}{I}$ and $e = \frac{b}{A}$ . Dissimilarity is the sum of turnover component and nestedness component. $\Delta AIC$ are calculated by subtracting the AIC of the null model (GLM with only one intercept) from the AIC of the fitted model. AUC are calculated with a Mann-Whitney U statistic.

## DISCUSSION

### Theoretical results

We derived a set of novel predictions of beta diversity patterns from the island biogeography theory. Under the classic MacArthur and Wilson framework, pairwise beta diversity patterns are determined by three factors: the size of the mainland species pool, extinction rates and colonization rates. Conditional on at least one island having species (*N*>0), only the expected Jaccard dissimilarity is independent of the size of mainland species pool. When conditioning on both islands having non-zero richness (*i+k* > 0 and *j+k* > 0), all indices are dependent on the size of mainland species pool. Ignoring empty islands has a huge influence on beta diversity patterns when the size of mainland species pool is small (e.g. less than 10 species), but the effect becomes negligible toward larger mainland species pool sizes, because of the rapidly decreasing probability of empty islands (Fig. 1a). This result stresses the need to include empty islands in empirical tests of the island biogeography theory (Wang et al. 2016).

Pairwise Jaccard dissimilarity increases with relative extinction rates (the ratio between extinction rates and colonization rates). This is because when extinction events become more frequent the chance of forming “checkerboard” patterns (Diamond 1975) grows, as shown by the increasing turnover component (Fig. 1b). The nestedness component first increases then decreases with relative extinction rates (Fig. 1c). The maximum nestedness is achieved when extinction rates are less than colonization rates (Fig. 1c). When relative extinction rates or the size of mainland species pool gets larger, the increase of Jaccard dissimilarity is mainly driven by the increasing turnover component (Fig. 1d). Indices in Podani’s family (the replacement and richness difference components) have similar qualitative behaviors, consistent with previous finding that the partitioned components in Baslega’s family are correlated with the partitioned components in Podani’s family (Legendre 2014; Baselga and Leprieur 2015). Island asymmetry, which takes into account the effect of isolation and area, does not change the qualitative behaviors of the indices (Fig 2), but increases the level of nestedness, because species in a small (distant) island are more likely to be a subset of species in a large (near) island.

### Neutral theory and null models

Our theoretical results are derived from a species-level neutral theory. We show that both the neutral GLM and the non-neutral GLM are successful in predicting species richness (alpha diversity) of birds in the Thousand Islands Lake (Fig. 3i, m). But only the non-neutral model predicts the observed beta diversity patterns. The patterns of partitioned components further reveal that the neutral GLM fails to predict the observed pairwise Jaccard dissimilarity because it overestimates turnover and underestimates nestedness (Fig. 3j-l). Our analysis does not include biological factors that may cause species-level non-neutrality, but the presented framework is flexible to include such information if available.

Null models have become one of the most important tools in ecology. Despite their increasing popularity, it has also been recognized that their assumptions (e.g. the widely used random shuffling approaches) can have serious limitations, especially in the presence of multiple processes (Gotelli and Ulrich 2012; Pigot and Etienne 2015; Miller et al. 2017; O’Dwyer et al. 2017). In terms of beta diversity, the entangled links among alpha, beta and gamma diversity are known to reduce the statistical power of randomization tests and generate ambiguity in their interpretations (Chase et al. 2011; Kraft et al. 2011; Qian et al. 2012, 2013; Ulrich et al. 2017). Our results support the use of mechanistic null models such as those based on neutral theory (O’Dwyer et al. 2009; Rosindell et al. 2012), maximum entropy theory (Xiao et al. 2015, 2016; O’Dwyer et al. 2017), and incidence functions (Hanski et al. 1996; Helm et al. 2006; Hanski 2009) to improve upon random-shuffling null models in hypothesis testing. These mechanistic models could generate multiple diagnostic patterns and allows stronger test for ecological theories.

### Future directions

Our results are derived from a mainland-island model where the colonization and extinction events on islands are independent of one another. More realistic modifications should be considered in order to predict biodiversity changes in a real landscape (Hanski 2009). For example, environmental heterogeneity and local dispersal are found to be important drivers of beta diversity in experimental studies (Grainger and Gilbert 2016; Gianuca et al. 2017; Ojima and Jiang 2017; Rodrigues and Diniz-Filho 2017). Moreover, in oceanic islands, high endemism arises from speciation events (Chen and He 2009; Gascuel et al. 2016; Steinbauer et al. 2016), which are likely to increase the level of species turnover. Future studies should also extend presence-based measures to abundance-based beta diversity measures which may provide additional insights into the dynamics of meta-community (Bashan et al. 2016; Kalyuzhny and Shnerb 2017). We believe that the integration of beta diversity patterns into the Theory of Island Biogeography offers new opportunities to infer community assembly processes.

## ACKNOWLEDGEMENTS

We thank Dr. James Rosindell for providing helpful comments for earlier drafts. We thank members in Jetz Lab and Vasseur Lab for feedbacks and dicussions.

## AUTHORSHIP

ML and DV conceived of the idea. The idea was further developed with DV and WJ’s input. ML conducted the analyses and wrote the manuscript. All authors assisted with revisions.

